# Mechanism of Actin Filament Severing and Capping by Gelsolin

**DOI:** 10.1101/2024.09.10.612341

**Authors:** Kyle R. Barrie, Grzegorz Rebowski, Roberto Dominguez

## Abstract

Gelsolin is the prototypical member of a family of Ca^2+^-dependent F-actin severing and capping proteins. A structure of Ca^2+^-bound full-length gelsolin at the barbed end shows domains G1G6 and the inter-domain linkers wrapping around F-actin. Another structure shows domains G1G3, a fragment produced during apoptosis, on both sides of F-actin. Conformational changes that trigger severing occur on one side of F-actin with G1G6 and on both sides with G1G3. Gelsolin remains bound after severing, blocking subunit exchange.

## Main text

Actin, the most abundant cytosolic protein in eukaryotes, exists in regulated equilibrium between monomeric (G-actin) and filamentous (F-actin) forms ^1^. F-actin networks drive essential cellular processes, including motility, cytokinesis, organelle trafficking, and endocytosis. Two types of proteins govern the dynamic turnover of actin networks: those promoting filament nucleation and elongation and those facilitating filament severing and depolymerization. Nucleators and elongators control the time and location for *de novo* F-actin polymerization whereas disassembly factors recycle the G-actin pool for new rounds of polymerization. F-actin disassembly factors belong to two major families: the ADF/cofilin-family, responsible for phosphorylation-regulated F-actin severing and pointed end depolymerization in conjunction with cyclase-associated protein ^2–4^, and the gelsolin-family, involved in Ca^2+^-dependent F-actin severing and barbed end capping ^5^. Members of the gelsolin family, including eight proteins in humans, have three to six copies of the gelsolin domain, consisting of a β-sheet sandwiched between two perpendicular α-helices, one long and one short (Extended Data Fig. 1 illustrates structures and nomenclatures used in gelsolin and actin). Gelsolin, the prototypical member of this family, has six gelsolin domains (G1 to G6). Secreted (or plasma) gelsolin features 24 extra amino acids at the N-terminus and is a major component of the actin scavenger system responsible for the clearance of F-actin released into circulation during tissue injury ^6^.

A crystal structure of Ca^2+^-free, inactive gelsolin reveals a compact arrangement of the six domains, forming a globule with G6 at its center (Extended Data Fig. 1b) ^7^. In this conformation, the actin-binding sites are mostly concealed. Crystal structures have also been determined of Ca^2+^-bound G1G3 and G4G6 in complex with G-actin ^8,9^. These structures reveal how Ca^2+^ binding releases autoinhibitory interactions, enabling the domains to open and bind actin. Gelsolin has two types of Ca^2+^-binding sites (Fig. 1a and Extended Data Fig. 1a-c); each domain has a type-2 Ca^2+^-binding site, and two additional type-1 Ca^2+^-binding sites are formed upon binding to actin, at the interfaces of actin-G1 and actin-G4. These structures, however, do not show how full-length gelsolin binds F-actin or how the six domains coordinate their interactions to produce severing and capping. Moreover, the G-to-F-actin transition involves a conformational change that transforms G-actin, described as having a “twisted” conformation, into the “flat” conformation of subunits in F-actin (Extended Data Fig. 1e) ^10^. The G1G3 and G4G6 structures show the twisted, G-actin conformation, not the flat conformation gelsolin targets when binding to F-actin.

**Fig. 1.**
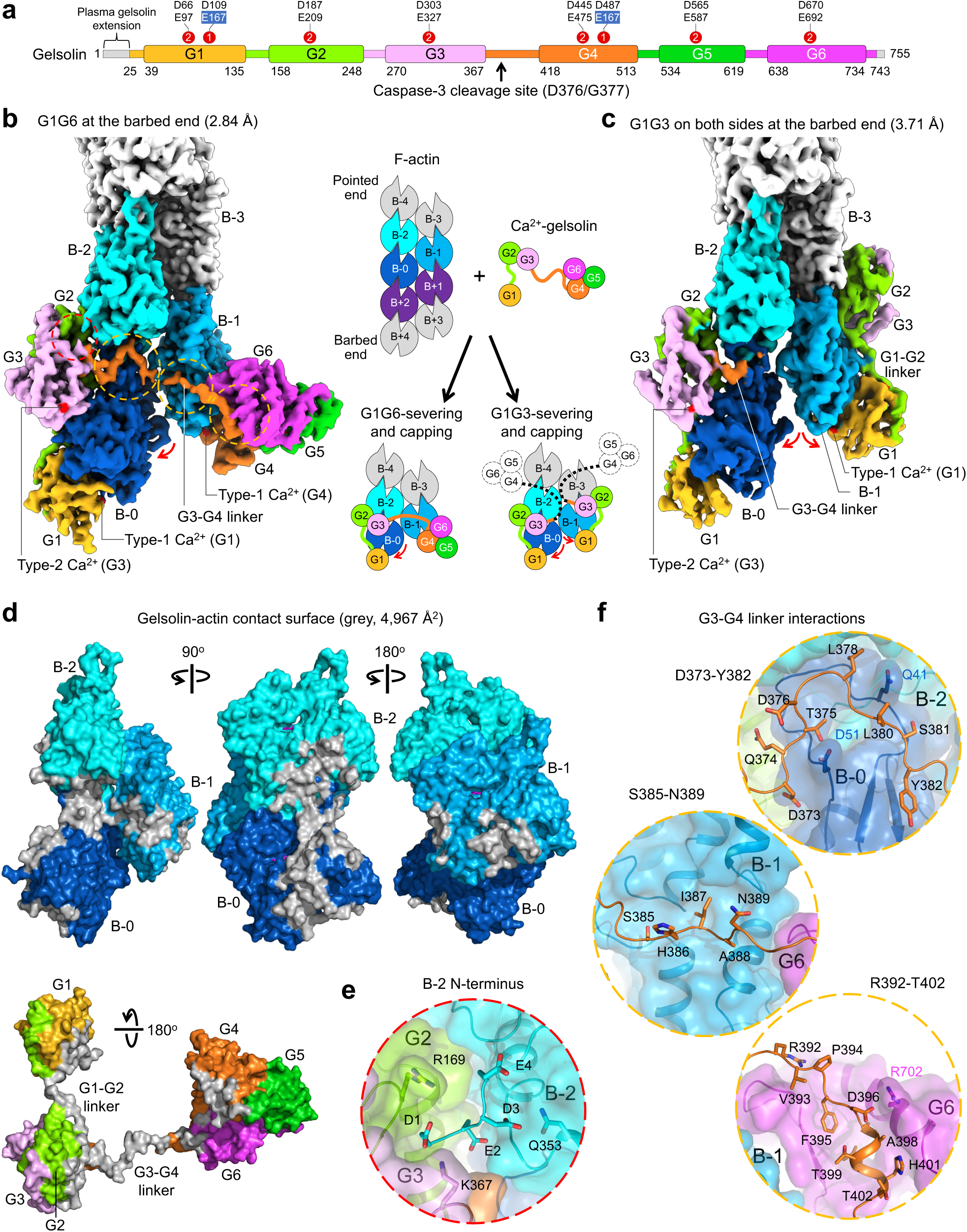
Structures of gelsolin at the barbed end. **a**, Domain organization of cytoplasmic and secreted (or plasma) gelsolin featuring 24 extra amino acids at the N-terminus), indicating the caspace-3 cleavage site during apoptosis and types1 and 2 Ca^2+^-binding sites with their main coordinating residues (actin residues shown in blue background). **b**-**c**, Cryo-EM maps of gelsolin G1G6 (full-length) and G1G3 at the barbed end. Gelsolin domains are colored according to part a, and actin subunits interacting with G1G6 are colored in different shades of blue. A cartoon representation of the severing reaction and resulting structures is shown in the middle. **d**, Contact surface area (4,967 Å^2^, grey) of the complex shown on F-actin (three different orientations) and gelsolin. **e**, Close-up view of B-2’s N-terminus stabilized by interactions with G2, G3, and B-2 itself (corresponding to the dotted red circle in part b). **f**, Close-up views of three regions of the G3-G4 linker (corresponding to dotted orange circles in part b). These regions interact, respectively, with B-0’s D-loop, a cleft between subdomains 3 and 4 of B-1, and G6.

### Gelsolin structures at the barbed end

Employing a strategy similar to the one we used to determine structures of the ends of F-actin ^11^, we obtained here cryo-electron microscopy (cryo-EM) structures of Ca^2+^-bound gelsolin at the barbed end. One structure, at 2.84-Å resolution, reveals gelsolin domains G1G3 and G4G6 interacting with the two long-pitch helices on opposite sides of F-actin (Fig. 1b, Extended Data Figs. 2-4 and Table 1). Gelsolin contacts three actin subunits, named B-0, B-1, B-2, and is predicted to interact with two more subunits (B+1 and B+2) before severing. The cryo-EM map shows density consistent with all eight Ca^2+^-binding sites of gelsolin being occupied, and actin subunits having Ca^2+^-ADP bound in their nucleotide cleft (Extended Data Fig. 5). Another structure, at 3.71-Å resolution and representing 13% of the gelsolin-containing particles, shows G1G3 bound on both sides of F-actin at the barbed end (Fig. 1c, Extended Data Figs. 6-7 and Table 1). In this structure, G4G6 was not visualized and likely does not interact with F-actin. Importantly, gelsolin was never observed bound in the middle of F-actin, suggesting severing occurs rapidly upon binding.

**Table 1.**
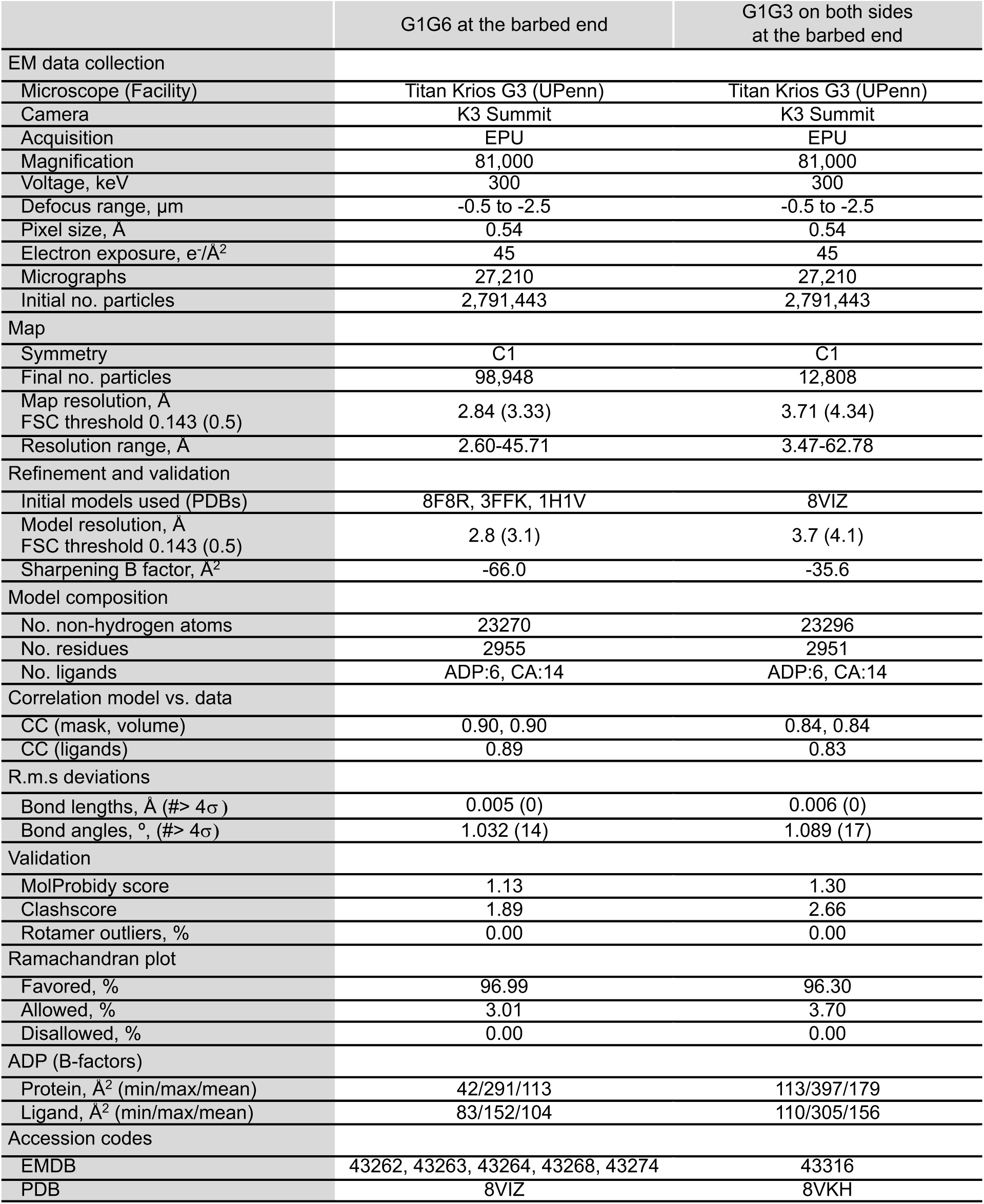
Cryo-EM data collection, processing, and model statistics.

### Interactions of G1G3

G1G3 contacts B-0 and B-2 along one of the long-pitch helices (Fig. 1d and Extended Data Figs. 8 and 9 illustrate the interactions and contact surface areas of gelsolin domain with actin subunits). The interaction of G1G3 with B-0 resembles that observed in the crystal structure with G-actin ^8^, with notable differences arising from the G-to-F-actin transition and newly observed contacts with B-2. G1 inserts its long helix into the hydrophobic cleft of B-0. The extended G1-G2 linker runs along the surface of B-0 toward B-2. This linker features a motif (^149^FKHV^152^) that is related to and binds the same site on actin as the LKKV motif of the WH2 domain ^12,13^. G2 interacts with B-0 and B-2. Its binding surface on B-2 resembles that of G1 on B-0, with one important distinction: the hydrophobic cleft of B-2 is occupied by the D-loop of B-0, preventing G2 from inserting its long helix into this cleft. Instead, the long helix of G2 lies on top of subdomain-2 of B-0 and the hydrophobic cleft of B-2. The G2-G3 linker is mostly folded as an α-helix and does not contact F-actin. G3 only makes a relatively minor contact with subdomain-1 of B-0. While actin’s N-terminus is disordered in most structures, that of B-2 (^1^DEDE^4^) binds in a pocket at the interface of G2, G3, and B-2 itself, where it is stabilized through interactions with R169 (G2), K367 (G3), and Q353 (B-2) (Fig. 1e). Before severing, contacts with B+2 should only involve G1. Everything described here applies to the binding of G1G3 on both sides of F-actin, where G1G3 additionally interacts with subunits B-1 and B-3, as well as B+1 before severing (Fig. 1c).

### Interactions of the G3-G4 linker

The G3-G4 linker, missing in previous G-actin-bound structures ^8,9^, spans the ∼92 Å distance separating G3 from G4 on opposite sides of F-actin. This extended conformation contrasts with the relatively compact conformation of this linker in inactive gelsolin ^7^, where it mediates autoinhibitory interactions between domains (Extended Data Fig. 1b). Because this linker is presumed flexible, it could potentially accommodate alternative positions of G1G3 relative to G4G6 on F-actin. However, the linker makes specific contacts with B-0, B-1, B-2, and G6 that define where on F-actin G4G6 binds relative to G1G3 (Fig. 1f). The first part of the linker (D373-Y382) forms a loop that lies on top of the D-loop of B-0. After crossing the crevasse between the two long-pitch helices, residues S385-N389 bind along a cleft formed between subdomains 3 and 4 of B-1. The following residues, R392-T402, interact with G6, before reaching G4 at the barbed end of B-1. The interaction with G6 is crucial, as it helps position G4G6 on the opposite long-pitch helix from G1G3, enabling gelsolin to form a pincer-like structure around F-actin. Thus, the G3-G4 linker plays a dual role, mediating autoinhibitory interactions in Ca^2+^-free gelsolin and coordinating the binding of the two gelsolin halves around F-actin in the Ca^2+^-bound state.

### Interactions of G4G6

Domains G4G6 form a clover-like ensemble and contact only B-1 (Fig. 1b,d and Extended Data Figs. 8 and 9). A similar arrangement was observed in the structure of G4G6 with G-actin ^8^, except in the current structure B-1 adopts a flat conformation. The interaction of G4 with B-1 resembles that of G1 with B-0, with G4’s long helix inserting into the hydrophobic cleft of B-1. G5 and the G4-G5 and G5-G6 linkers do not contact B-1, whereas G6 contacts only a small surface on subdomain-3 of B-1. Before severing, contacts with B+1 should only involve G4.

### Conformational changes in F-actin

The three subunits gelsolin contacts at the barbed end have the flat, F-actin conformation. Compared to subunits in F-actin, B-2 remains mostly unchanged, whereas B-1 remains in the same position but undergoes changes in the hydrophobic cleft (Fig. 2a-c). These changes affect C-terminal residues I345-F375 and the W-loop on each side of the hydrophobic cleft, which in F-actin hosts the D-loop of B+1 but in the current structure interacts with the long helix of G4. Similar changes, albeit more pronounced, are observed in B-0, whose hydrophobic cleft hosts the long helix of G1 (Fig. 2d). The D-loop of B-0 is also slightly shifted due to its interaction with the G3-G4 linker. Yet, the most striking change is a ∼6° rotation of B-0 away from the main F-actin axis, which displaces residues at the barbed end by up to 5.5 Å (Fig. 2e,f). Because the fulcrum of this rotation falls near the D-loop, it is most likely the binding of G2G3 and surrounding linkers that causes it. Consistently, severing is observed with a hybrid construct in which the G2G3 domains of gelsolin replace those of CapG, a three-domain member of this family lacking severing activity ^14^. Moreover, when G1G3 is bound on both sides of F-actin, B-0 and B-1 are both rotated (Extended Data Fig. 10), whereas subunits at the free barbed end are not ^11^. Additionally, in the G1G3 structure the hydrophobic cleft of B-1 undergoes large changes, similar to those of B-0, suggesting that G1 binding produces these changes.

**Fig. 2.**
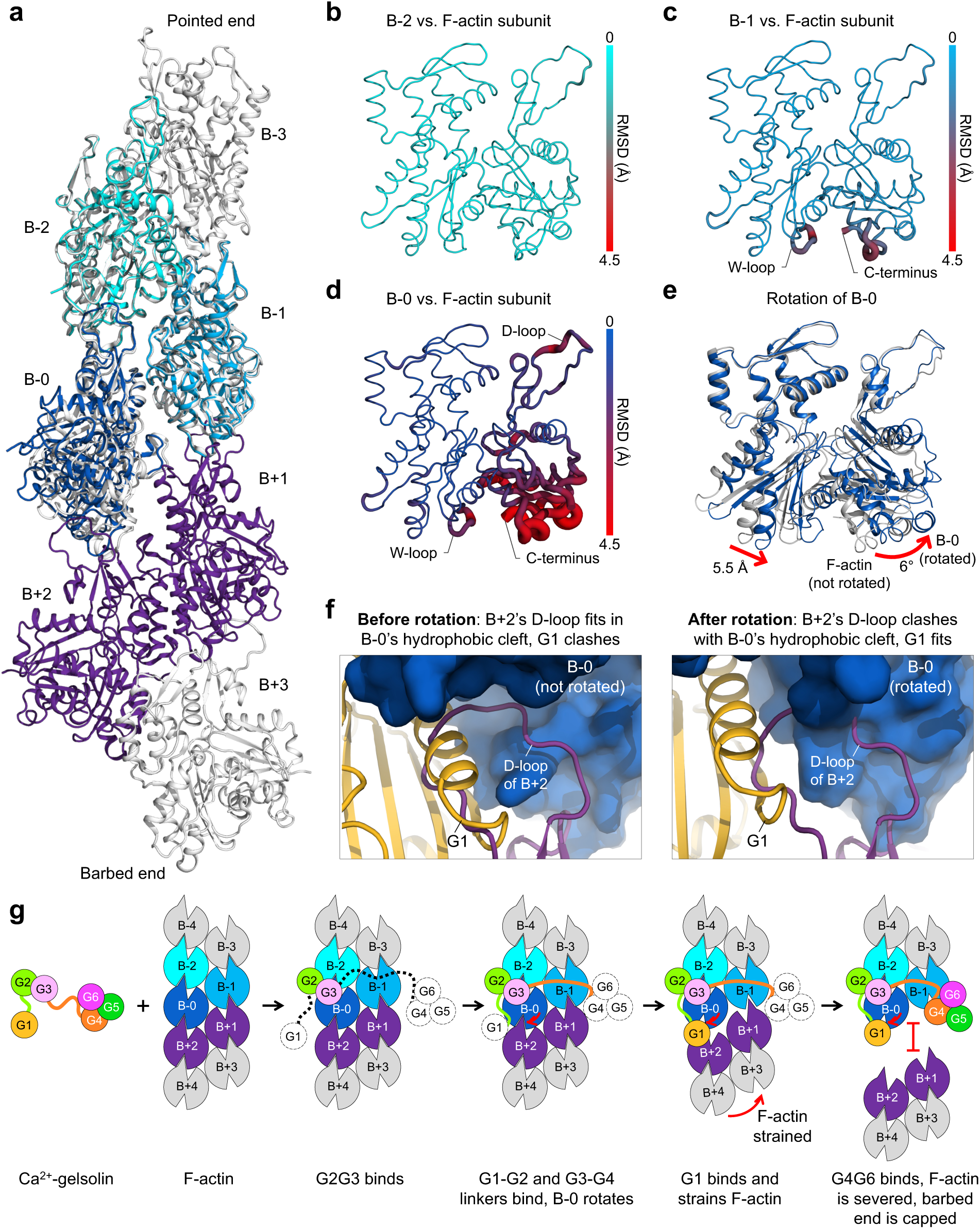
Conformational changes in actin subunits and severing model. **a**, Actin subunits B-0, B-1, and B-2 (colored in different shades of blue) from the G1G6-bound structure superimposed onto F-actin (PDB: 8F8P). F-actin subunits are colored grey, except those predicted to interact with gelsolin before severing, which are colored purple. **b**-**d**, Cartoon putty representation of B-2, B-1, and B-0 colored and rendered according to RMSD from the corresponding F-actin subunit (as indicated by side bars). B-2 remains mostly unchanged, whereas the hydrophobic clefts of B-1 and especially B-0 change substantially due to the binding of G4 and G1, respectively. The D-loop of B-0 also changes due to interactions with the G3-G4 linker. **e**, B-0 (blue) is rotated by 6° compared to its corresponding subunit in F-actin (grey), with residues at the barbed end moving by up to 5.5 Å (this rotation occurs in addition to the changes shown in part d). **f**, Comparison of the position of G1 on F-actin before and after the rotation of B-0. In F-actin, B+2’s D-loop fits in B-0’s hydrophobic cleft and G1 clashes. After rotation, G1 fits in B-0’s hydrophobic cleft and B+2’s D-loop clashes. **g**, Severing and capping model. Major steps, described below each diagram and in the main text, are also illustrated in Supplementary Video 1.

### Mechanism of severing and capping

Since its discovery in 1979 ^15^, gelsolin has been intensively investigated ^5^. The accumulated knowledge and current structures allow us to propose a severing model. G1 and G4G6 bind G-actin in a Ca^2+^-independent and Ca^2+^-dependent manner, respectively, but neither fragment binds F-actin ^16–18^. F-actin binding requires G2G3, which by itself does not sever ^16–18^. The minimal severing fragment contains G1 plus an F-actin-binding element, which can be the G1-G2 linker ^19^ or even an unrelated F-actin-binding domain from α-actinin ^20^. While early studies suggested G1G3 harbored most of the severing activity, it later emerged that this was due to G1G3 binding cooperatively on both sides of F-actin to compensate for the absence of G4G6 ^21^, emphasizing the importance of all the gelsolin domains for efficient severing. The cryo-EM structures rationalize these findings and support a multi-step severing model (Fig. 2g and Supplementary Video 1). First, G2G3 binds at the interface between B-0 and B-2. Second, the G1-G2 and G3-G4 linkers bind. These interactions trigger the third and fourths steps, which likely occur simultaneously. Third, G1 and G4 are positioned near their binding sites on opposite sides of F-actin but cannot bind due to steric clashes of their long helices with the D-loops of B+1 and B+2. Fourth, B-0 is rotated away from the filament axis. Fifth, the rotation of B-0 disrupts its interaction with the D-loop of B+2 while favoring binding of G1 in the hydrophobic cleft of B-0 (Fig. 2f), considered the highest affinity gelsolin-actin contact ^18^. Sixth, the binding of G1 drives a wedge between subunits B-0 and B+2 of the long-pitch helix, which strains the filament and allows G4 to bind at the barbed end of B-1. Seventh, the coordinated pincer motion of G1 and G4G6 on both sides of F-actin leads to severing. Capping is a consequence of severing, since after severing G1 and G4 occupy the hydrophobic clefts of subunits at the barbed end, which can no longer accept the D-loop of incoming subunits. Subunit dissociation after severing is also inhibited through stabilization of the B-0/B-2 longitudinal contact by interactions with G2G3 and surrounding linkers.

While this model shares features with previous models ^5,9,22–24^, the relative position of gelsolin domains on F-actin, the essential role of the G3-G4 linker, and the rotation of B-0 are important elements of our model that remained unresolved. Another surprise was to observe G1G3 bound on both sides at the barbed end but never in the middle of F-actin, which strongly supports the cooperative severing model proposed for this fragment ^21^. Importantly, the G1G3 fragment is physiologically generated by caspase-3 during apoptosis, which leads to Ca^2+^-independent F-actin severing, cell rounding and nuclear fragmentation ^25^.

## Methods

### Proteins

Actin was purified from rabbit skeletal muscle ^26^. The cDNA coding for full-length human cytoplasmic gelsolin was obtained from Tatyana Svitkina and PCR-amplified with primers designed to add an N-terminal 6x histidine tag and a C-terminal strep tag (forward: TTCGAATTCATGCATCACCATCATCACCACATGGTGGTGGAACACCCCGA, reverse: TTCGTCGACTCACTTTTCGAACTGTGGATGACTCCAGGCAGCCAGCTCAGCCAT). The PCR product was cloned between the EcoRI and SalI sites of vector pRSF1 (Novagen). The protein was expressed in ArcticExpress (DE3) RIL cells (Agilent Technologies) grown in Terrific Broth (TB) medium for 7 h at 37 °C to an optical density at 600 nm (OD_600_) of 1.2, followed by 24 h growth at 10 °C in the presence of 0.3 mM isopropyl-β-D-1-thiogalactoside (IPTG). Cells were harvested by centrifugation (50,000 × *g* for 20 min), resuspended in 20 mM HEPES pH 7.5, 300 mM NaCl, 2 mM EGTA, cOmplete protease inhibitor (Roche) and lysed using a Microfluidizer (Microfluidics). The protein was affinity-purified on a Ni-NTA column, followed by a Strep-Tactin column. The purified protein was concentrated to 6 mg/mL in 20 mM HEPES pH 7.5, 150 mM NaCl, 1 mM EGTA, and 40 mM D-biotin. Aliquots were flash frozen in liquid nitrogen and stored at −80 °C.

### Cryo-EM sample preparation, data acquisition, and processing

ATP-actin (50 µM) in G-actin buffer (2 mM Tris pH 8.0, 0.2 mM CaCl_2_, 0.2 mM ATP, 0.5 mM DTT) was added to a 50 µL solution containing 14 µM gelsolin in Ca^2+^-F-actin buffer (20 mM HEPES pH 7.5, 50 mM KCl, 1 mM CaCl_2_, 0.2 mM ATP, and 0.5 mM DTT) and incubated at room temperature for 2 hours. This combination produces many short filaments in cryo-EM micrographs, which is the ideal situation to study filament ends ^11^. For cryo-EM analysis, samples (3 µL) were applied onto glow-discharged (90 s, PELCO easiGlow) 200-mesh R1.2/1.3 Quantifoil holey carbon grids (Electron Microscopy Sciences). The grids were blotted for 2.5 s with force 7 using Whatman 41 filter paper and vitrified by plunging into liquid ethane using a Vitrobot Mark IV (ThermoFisher Scientific). Cryo-EM datasets were collected in five sessions using the EPU software (ThermoFisher Scientific) on a FEI Titan Krios transmission electron microscope operating at 300 kV and equipped with a Gatan K3 direct electron detector with an energy quantum filter. Images were collected with a defocus range of -0.5 to -2.5 µm, a total dose of 45 e^-^/Å^2^, and a nominal magnification of 81,000x in super-resolution mode, resulting in a pixel size of 0.54 Å.

Cryo-EM movies (27,210) were imported into CryoSPARC v4.0 ^27^ and binned by 2 during patch motion correction, resulting in a pixel size of 1.08 Å. After patch contrast transfer function (CTF) estimation, micrographs with a CTF fit > 8 Å were excluded, resulting in 26,860 selected micrographs. Particle picking was performed with the program Topaz ^28^ as we have described ^11^. Starting from a subset of ∼1,000 manually picked F-actin ends, multiple iterations of Topaz training, particle picking, and reference-free 2D classification resulted in a starting dataset of 2,791,443 particles. Particles were extracted with a box size of 416 pixels (449 Å) and binned by 8 to a box size of 52 pixels. Reference-free 2D classification was used to remove particles lacking structural features. Accepted particles were used to generate three ab-initio volumes, one resembling F-actin and two “junk” volumes. Particles belonging to the junk volumes were discarded, and the remaining particles were subjected to heterogeneous refinement using as input three identical F-actin ab-initio volumes. This procedure produced three classes, corresponding to the middle of F-actin, the free pointed end, and the gelsolin-bound barbed end. Free barbed ends (i.e., without gelsolin) were not observed. Moreover, despite using multiple strategies and attempts, particles featuring gelsolin bound to the middle of F-actin (i.e., before severing) were not observed. This state does not appear to exist in the dataset, consistent with gelsolin severing F-actin promptly after binding.

Particles in the gelsolin-bound barbed end class (482,293) were re-centered, such that the filament ends were at the center of the box and extracted using a box size of 416 pixels without binning. These particles were used as input for a new ab-initio reconstruction with one class, resulting in a map containing good density for G1G3 but relatively poor density for G4G6. Focused 3D classification without image alignment, using 8 classes and a mask encompassing G1G6, B-0, and B-1, further separated particles into classes corresponding to the middle of F-actin, poorly resolved ends, well-resolved ends, and well-resolved ends containing an additional actin subunit. Particles belonging to the middle of F-actin and poorly resolved ends were discarded. Particles belonging to the well-resolved ends containing an additional actin subunit were re-centered and re-extracted such that all the ends were in register. Particles from the re-centered and well-resolved stacks were combined and used in non-uniform refinement, resulting in a map containing clear density for G1G3 and G4G6. Particles lacking clear density for G5G6 were removed through two rounds of focused 3D classification without image alignment, first using a mask encompassing G4G6 and then a mask encompassing G5G6. The resulting final particle stack (98,948) was subjected to global and local CTF refinement followed by non-uniform refinement to produce the final “consensus” map at 2.84 Å resolution. To improve the density in regions of the structure that appear to move semi-independently, local refinements were performed using three different masks (encompassing G1G3, B-0, B-2; G4G6, B-1; B-3, B-4, B-5). The maps resulting from the local refinements were re-sampled, using the consensus map as reference, and combined to produce the final map used in model building and refinement.

The 98,948 particles used to obtain the final map of G1G6 at the barbed end were subtracted from the initial stack of 482,293 gelsolin-containing particles. The remaining stack (381,176 particles) was used as input for a new ab-initio reconstruction with one class. As described above, iterative 3D classification without image alignment, using masks focused on different regions of the map (Extended Data Fig. 6), converged on a 12,808-particle stack representing G1G3 bound on both sides of F-actin at the barbed end. These particles were subjected to local and global CTF refinement followed by non-uniform refinement to produce a consensus 3.71-Å resolution map of G1G3 at the barbed end that was used in model building and refinement.

Map quality was evaluated using programs 3DFSC ^29^ and cryoEF ^30^. 3DFSC yielded sphericity values of 0.976 and 0.750 for the G1G6 and G1G3 maps, respectively. The efficiency of the orientation distribution (E_od_) determined with cryoEF was 0.76 and 0.79 for the G1G6 and G1G3 maps, respectively. Local resolutions were calculated in CryoSPARC using a Fourier shell correlation (FSC) cutoff of 0.5 (Extended Data Figs. 3, 4, and 7).

### Model building and refinement

Coot ^31^ and Phenix ^32^ were used for model building and refinement, starting from a high-resolution cryo-EM structure of the free barbed end of F-actin (PDB code 8F8R) ^11^ and crystal structures of Ca^2+^-bound G1G3 and G4G6 in complex with G-actin (PDB codes: 3FFK and 1H1V) ^8,9^. Non-crystallographic symmetry constraints were not imposed on actin subunits during refinement. Individual atomic positions and temperature factors were refined for both structures using the real-space-refinement function of Phenix. Final model quality and refinement statistics are given in Table 1. Figures were prepared with the programs PyMOL (Schrödinger, LLC) and ChimeraX ^33^.

## Acknowledgements

Data collection was performed at The Beckman Center for Cryo-Electron Microscopy, University of Pennsylvania (Research Resource Identifier: SCR_022375). We thank Stefan Steimle for assistance with data collection and Tatyana Svitkina for providing gelsolin’s cDNA.

## Data availability

All maps and models generated have been deposited to the EMDB and PDB. G1G6 at the barbed end: consensus map (EMD-43262), local refinements (EMD-43263, EMD-43264, EMD-43268), combined map (EMD-43274), and coordinates (PDB: 8VIZ). G1G3 on both sides at the barbed end: consensus map (EMD-43316) and coordinates (PDB: 8VKH).

## Author Contributions

K.R.B. and R.D. conceived the study and designed the experiments. K.R.B. and G.R. prepared the proteins. K.R.B. performed cryo-EM experiments including sample and grid preparation, data collection, and data analysis. K.R.B and R.D. built and refined the atomic models. R.D. procured funding. K.R.B and R.D. wrote the paper and prepared figures.

---

**Extended Data Fig. 1.**
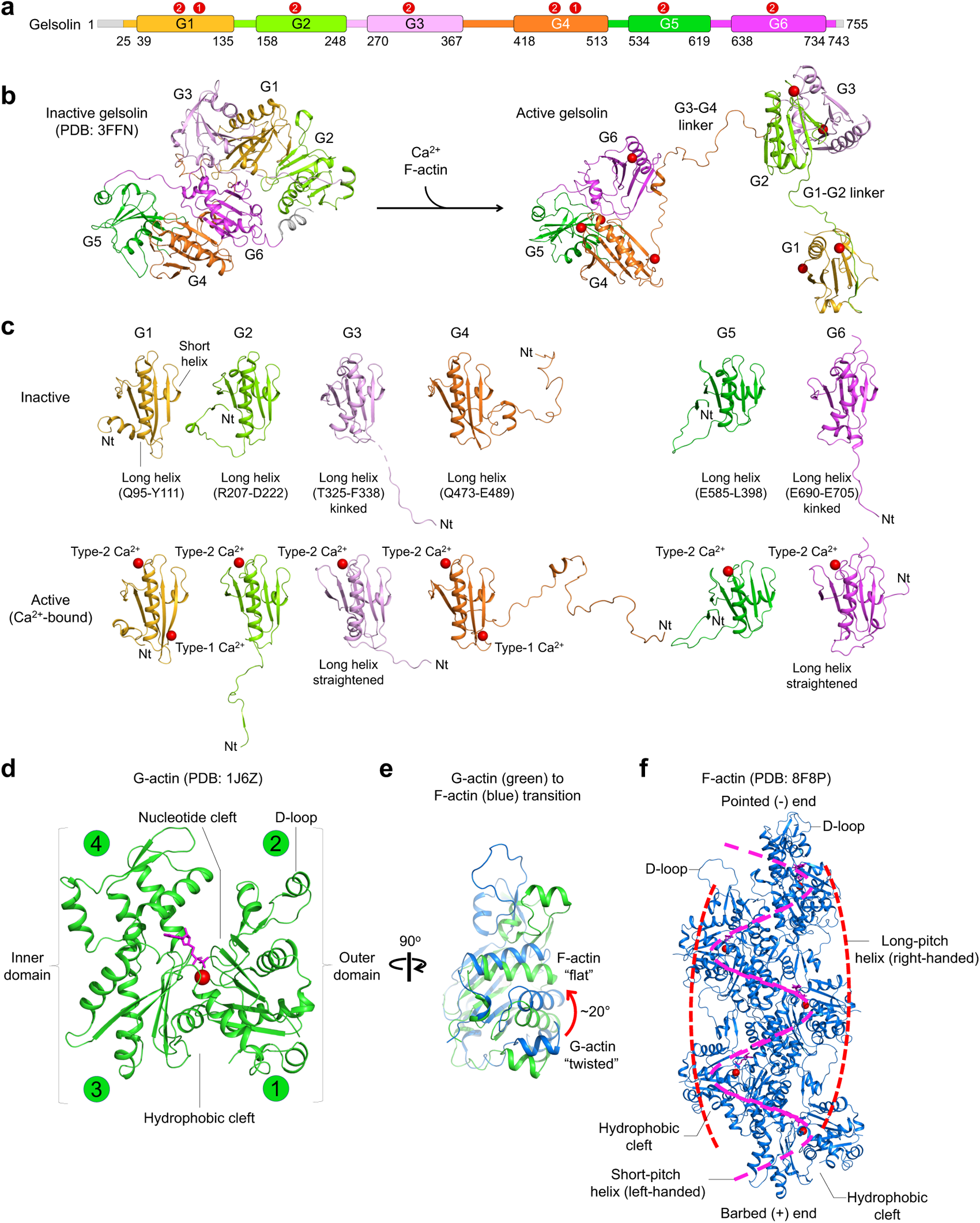
Structures and nomenclatures used in gelsolin and actin. **a**, Domain organization of gelsolin (as in Fig. 1a). **b**, Structures of full-length inactive gelsolin (PDB: 3FFN) and Ca^2+^-bound, active gelsolin from the complex with F-actin. **c**, Gelsolin domains and their preceding linkers (colored as in part a) in the inactive and active (Ca^2+^-bound, red) structures, indicating nomenclatures used in the main text. Note the changes in the position of the linkers and the straightening of the long helices of G3 and G6 from the inactive to the active structures. **d**, Structure of G-actin (PDB: 1J6Z), indicating nomenclatures used in the main text. Numbers in green circles indicate actin subdomains 1 to 4. **e**, G- to F-actin transition. When the inner domain is superimposed, the outer domain rotates by ∼20° from the twisted (G-actin, green) to the flat (F-actin, blue) conformation. **f**, Structure of F-actin (PDB: 8F8P), indicating nomenclatures used in the main text.

**Extended Data Fig. 2.**
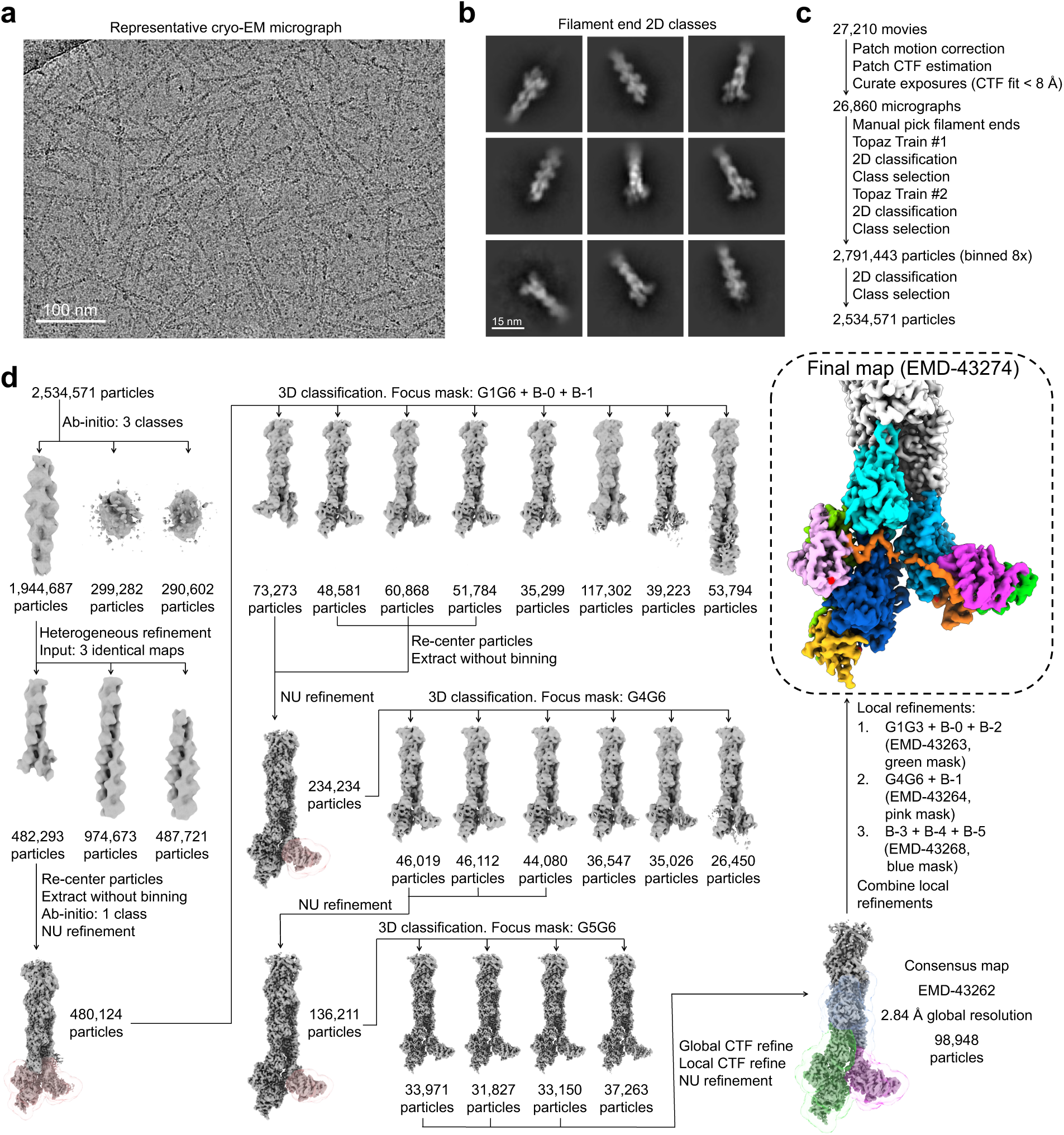
Cryo-EM workflow for the map of G1G6 at the barbed end. **a**, Representative micrograph. **b**, 2D class averages of F-actin ends. **c**, Cryo-EM movie processing and particle picking strategy. **d**, Particle processing workflow and masks (pink, blue, green, and magenta) used in focused 3D classification and local refinement, resulting in the final map at 2.84-Å resolution (see Methods).

**Extended Data Fig. 3.**
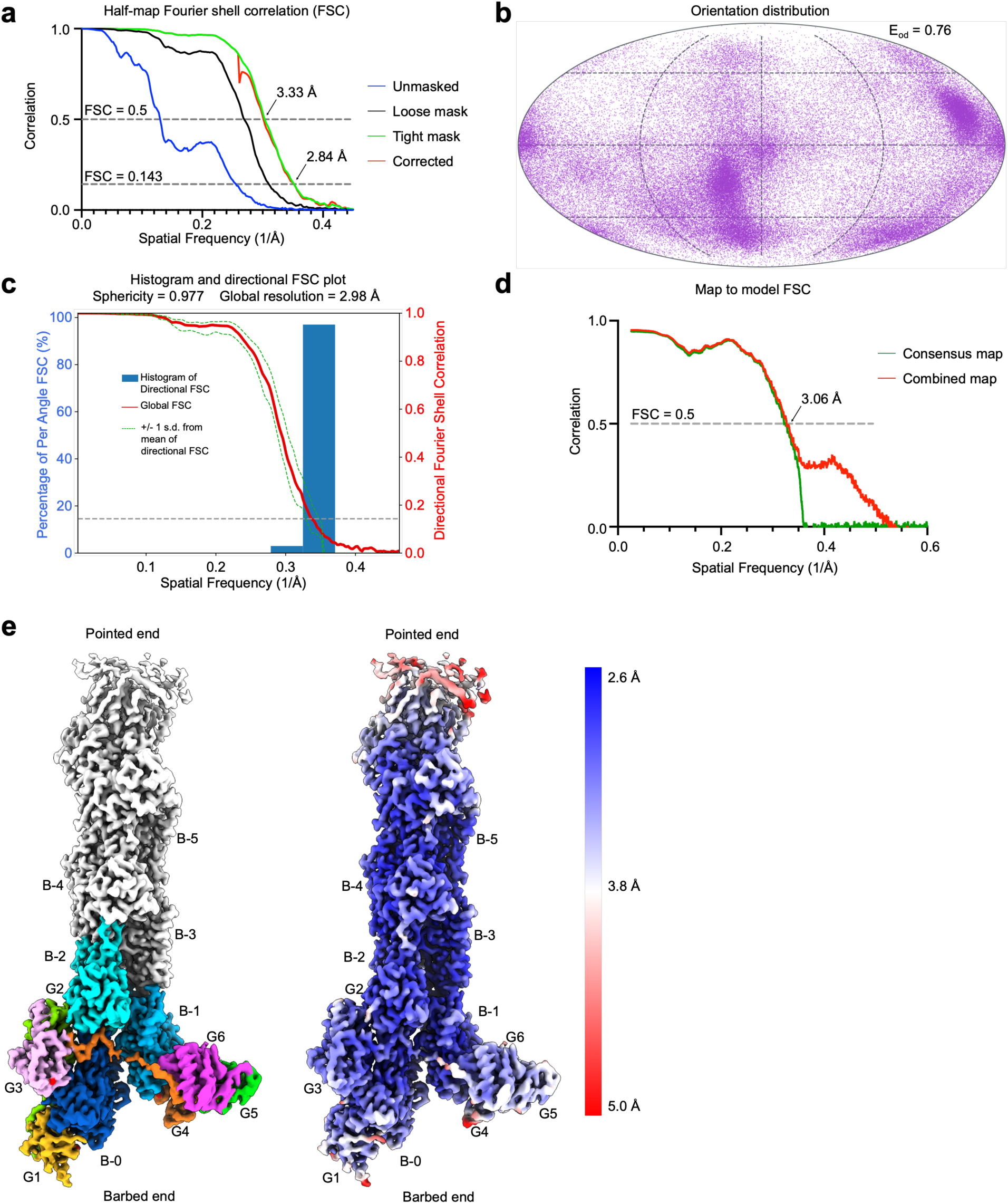
Quality and resolution assessment of the map of G1G6 at the barbed end. **a**, Half-map Fourier shell correlation (FSC) resolution analysis of the consensus map (2.84 Å at FSC = 0.143). **b**, Orientation distribution of particles used in the consensus map determined with the program cryoEF ^29^. The consensus map has a calculated efficiency (E_od_) of 0.76. **c**, 3D Fourier shell correlation calculated using 3DFSC ^28^. The consensus map has a sphericity of 0.977 and a global resolution of 2.98 Å. **d**, Map to model FSC determined with the program Phenix ^31^ for the consensus (green) and combined (red) maps. **e**, Cryo-EM structure colored by actin subunits and gelsolin domains (left) or by local resolution, as indicated by the scalebar (right). Actin subunits used in model refinement are labeled (B-0 to B-5).

**Extended Data Fig. 4.**
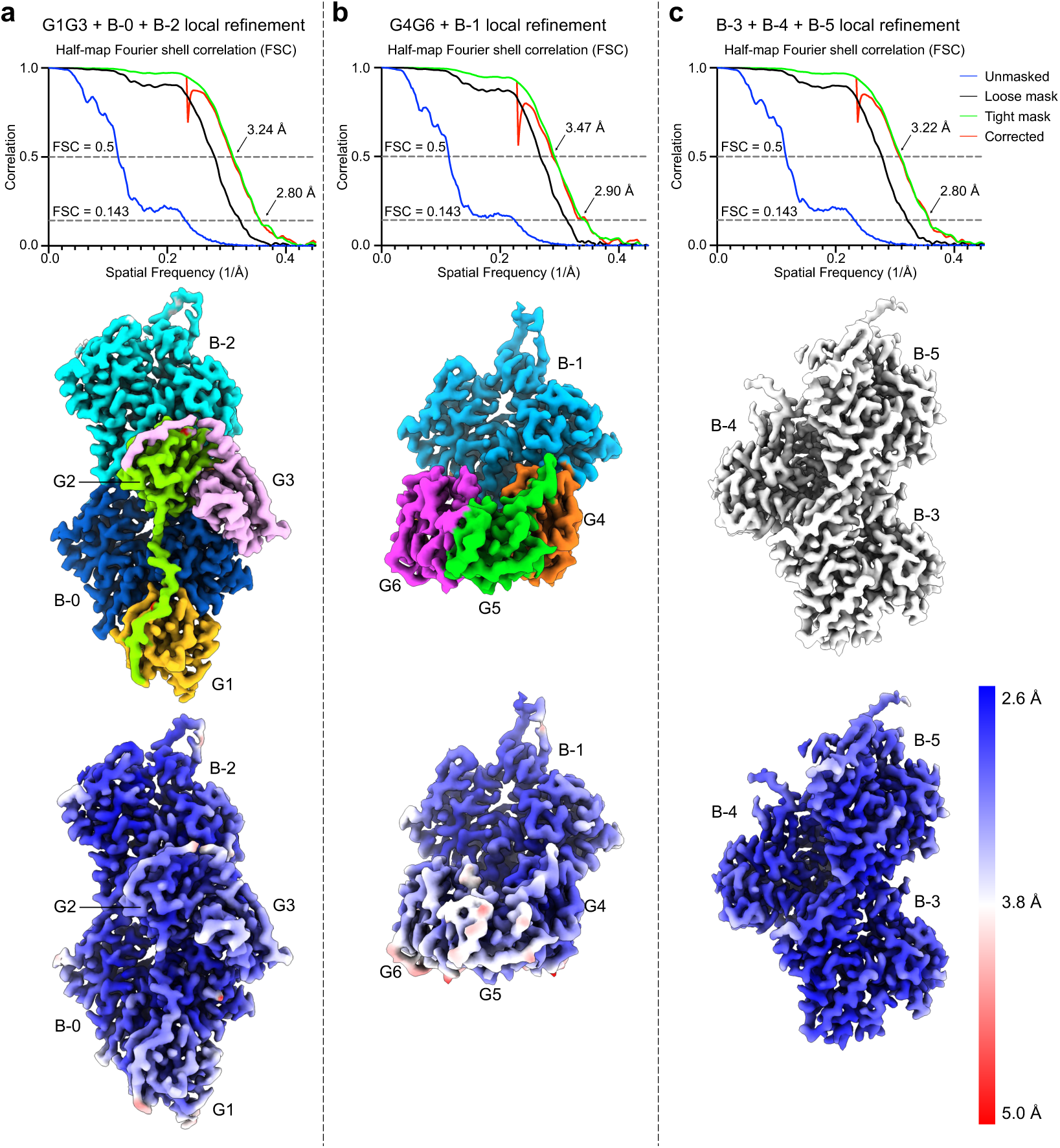
Resolution assessment of the local refinements leading to the combined map of G1G6 at the barbed end. **a**-**c**, FSC resolution analysis of the local-refined maps and local resolution estimates. Actin subunits and gelsolin domains included in each of the three masks are colored as in Fig. 1b (main text) or by local resolution, as indicated by the scalebar (right).

**Extended Data Fig. 5.**
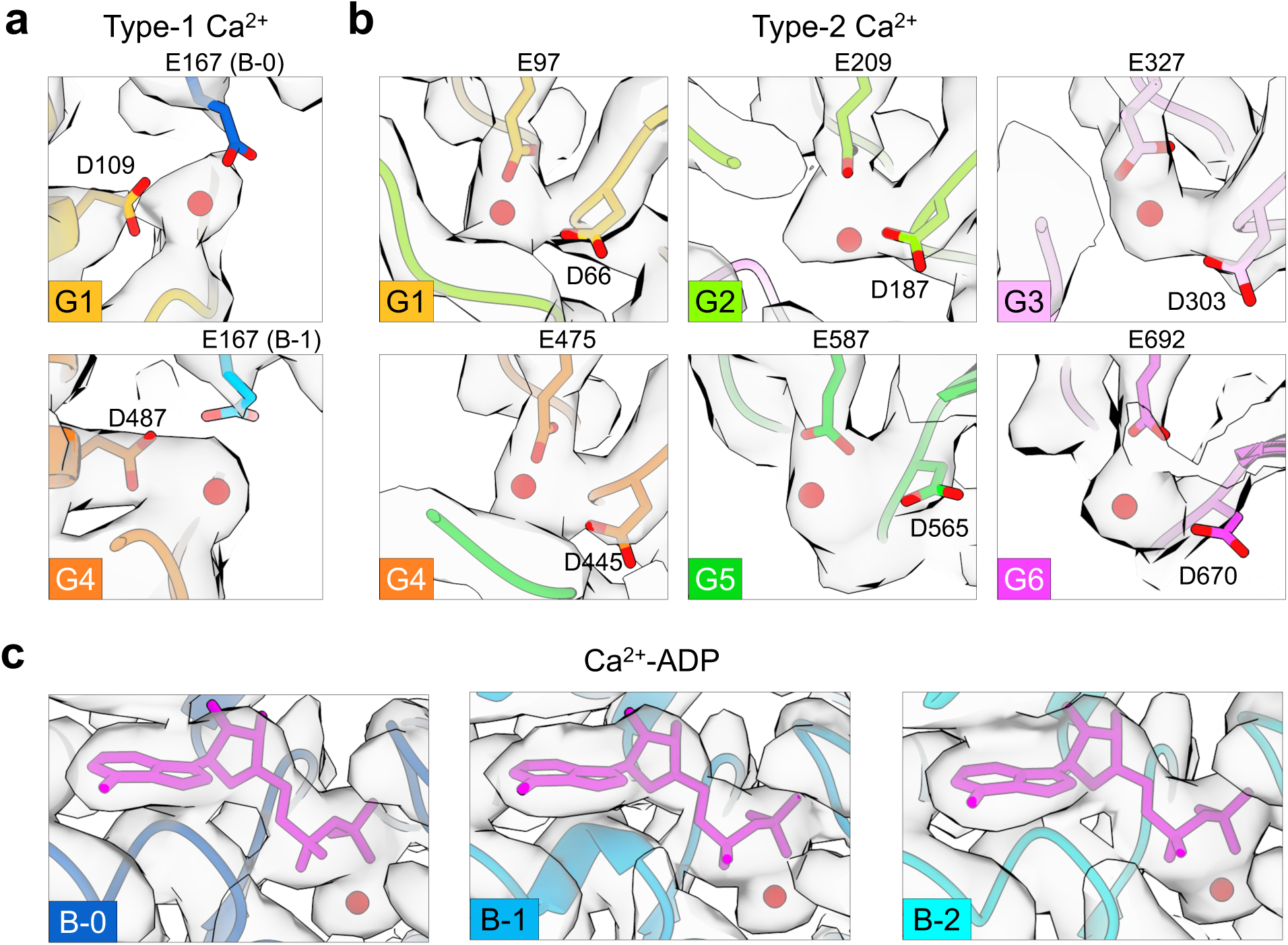
Cryo-EM map around the Ca^2+^ and Ca^2+^-ADP sites of gelsolin and actin subunits. **a**-**c**, All-atom representation of the final refined model around the Ca^2+^ and Ca^2+^-ADP binding sites of gelsolin (**a** and **b**) and actin subunits (**c**) and corresponding density in the combined cryo-EM map of G1G6 at the barbed end (atoms are colored according to Fig. 1b in the main text).

**Extended Data Fig. 6.**
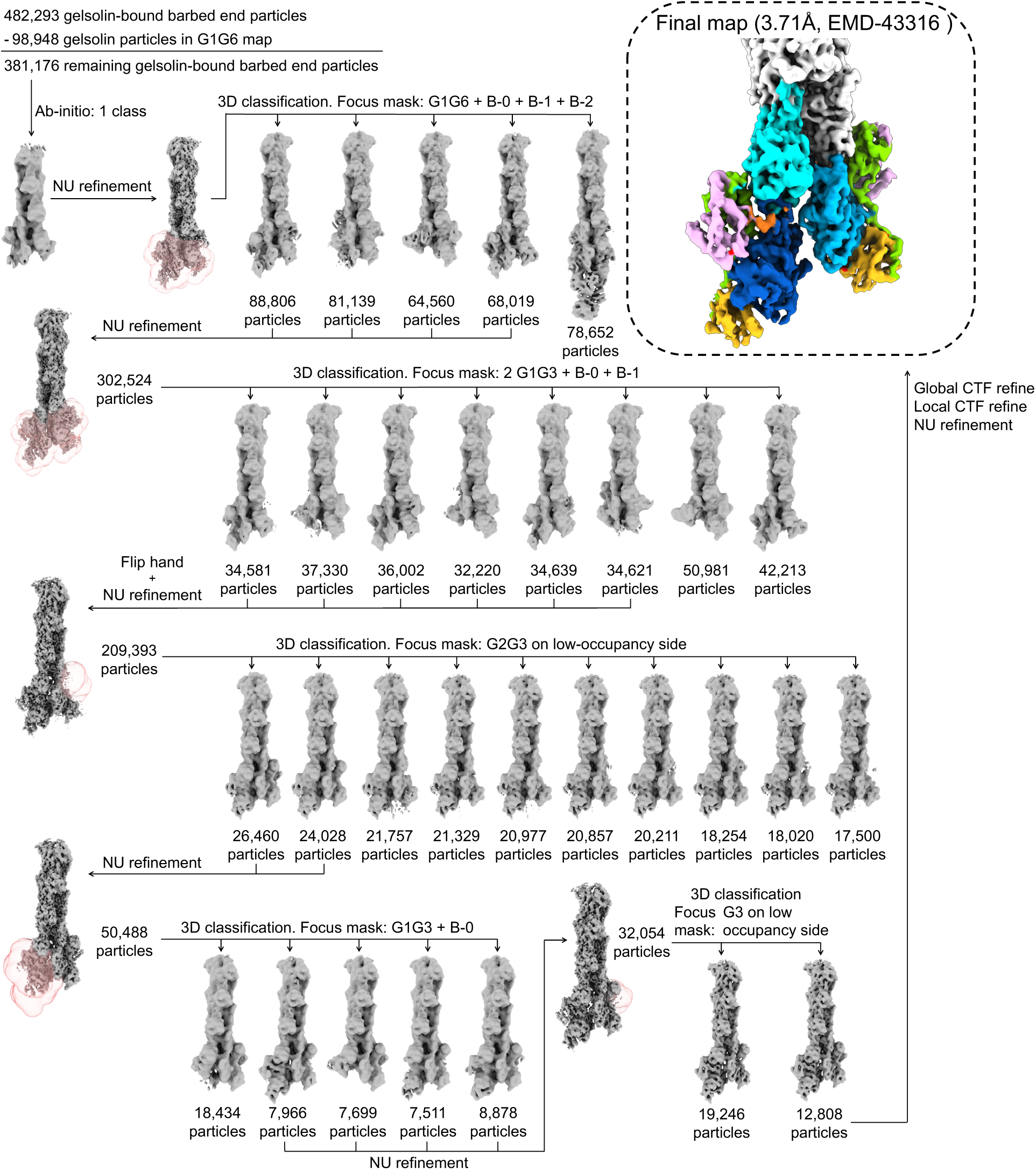
Cryo-EM workflow for the map of G1G3 at the barbed end. Particle processing workflow and masks (pink) used in focused 3D classification, resulting in the final map at 3.71-Å resolution (see Methods).

**Extended Data Fig. 7.**
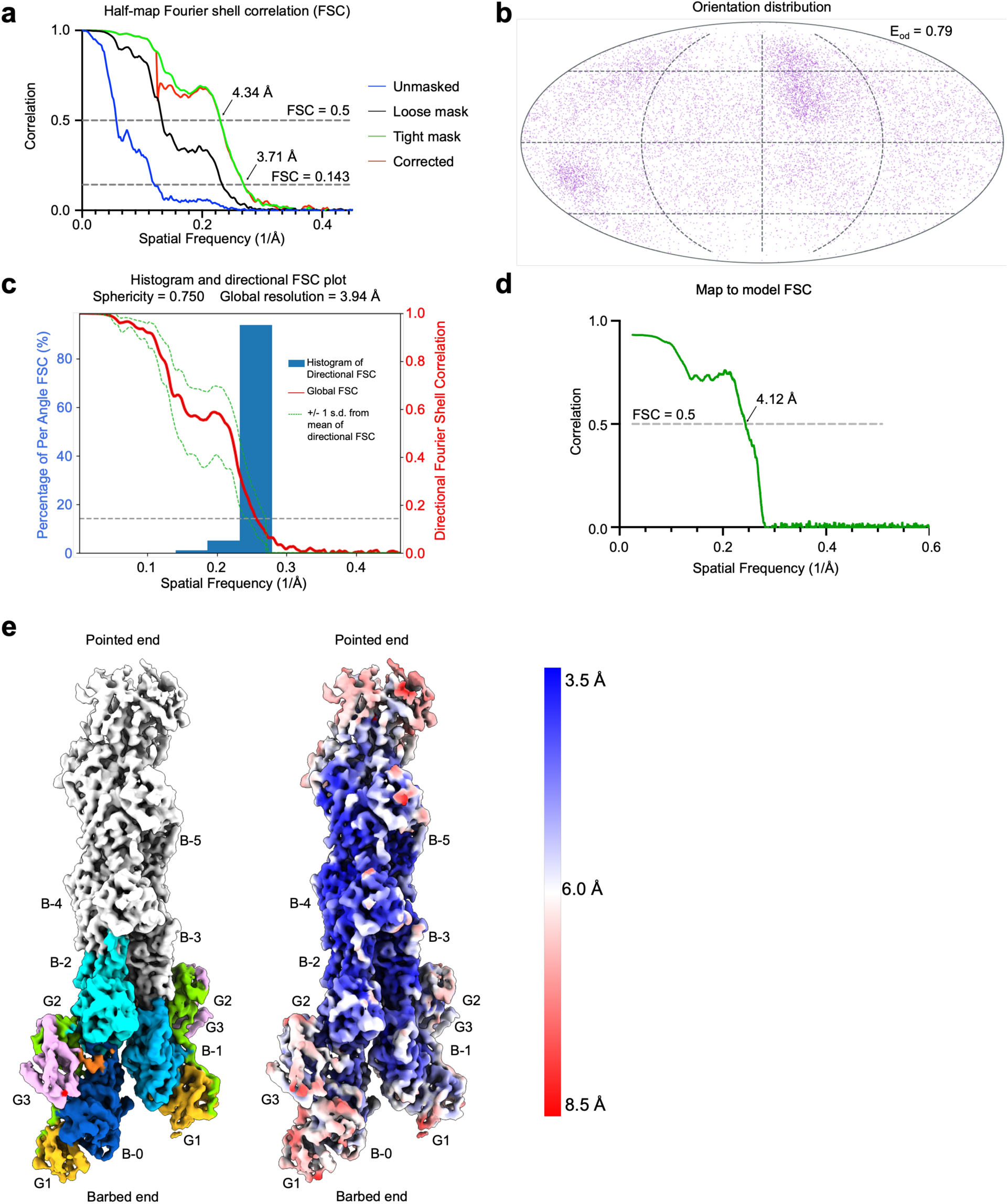
Quality and resolution assessment of the map of G1G3 at the barbed end. **a**, Half-map Fourier shell correlation (FSC) resolution analysis of the map (3.71 Å at FSC = 0.143). **b**, Orientation distribution of particles used in the map determined with the program cryoEF ^29^. The map has a calculated efficiency (E_od_) of 0.79. **c**, 3D Fourier shell correlation calculated using 3DFSC ^28^. The map has a sphericity of 0.75 and a global resolution of 3.94 Å. **d**, Map to model FSC determined with the program Phenix ^31^. **e**, Cryo-EM structure colored by actin subunits and gelsolin domains (left) or by local resolution, as indicated by the scalebar (right). Actin subunits used in model refinement are labeled (B-0 to B-5).

**Extended Data Fig. 8.**
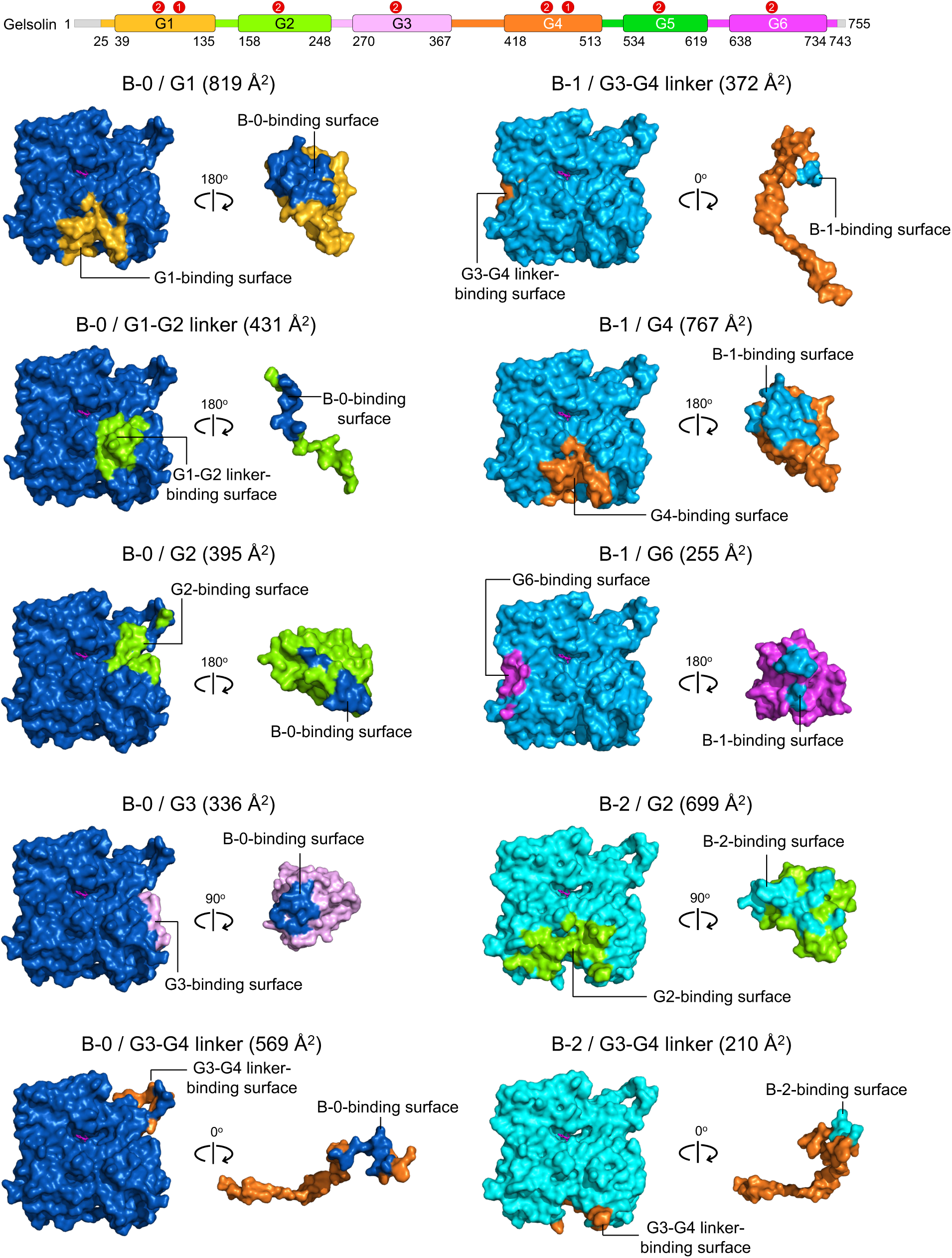
Contact surface areas between actin subunits and gelsolin domains and linkers. For each interaction, the contact area in the actin subunit is colored according to the gelsolin region it binds and *vice versa*. Contact surface areas (given for each interaction) were calculated using the server GetArea (https://curie.utmb.edu/getarea.html).

**Extended Data Fig. 9.**
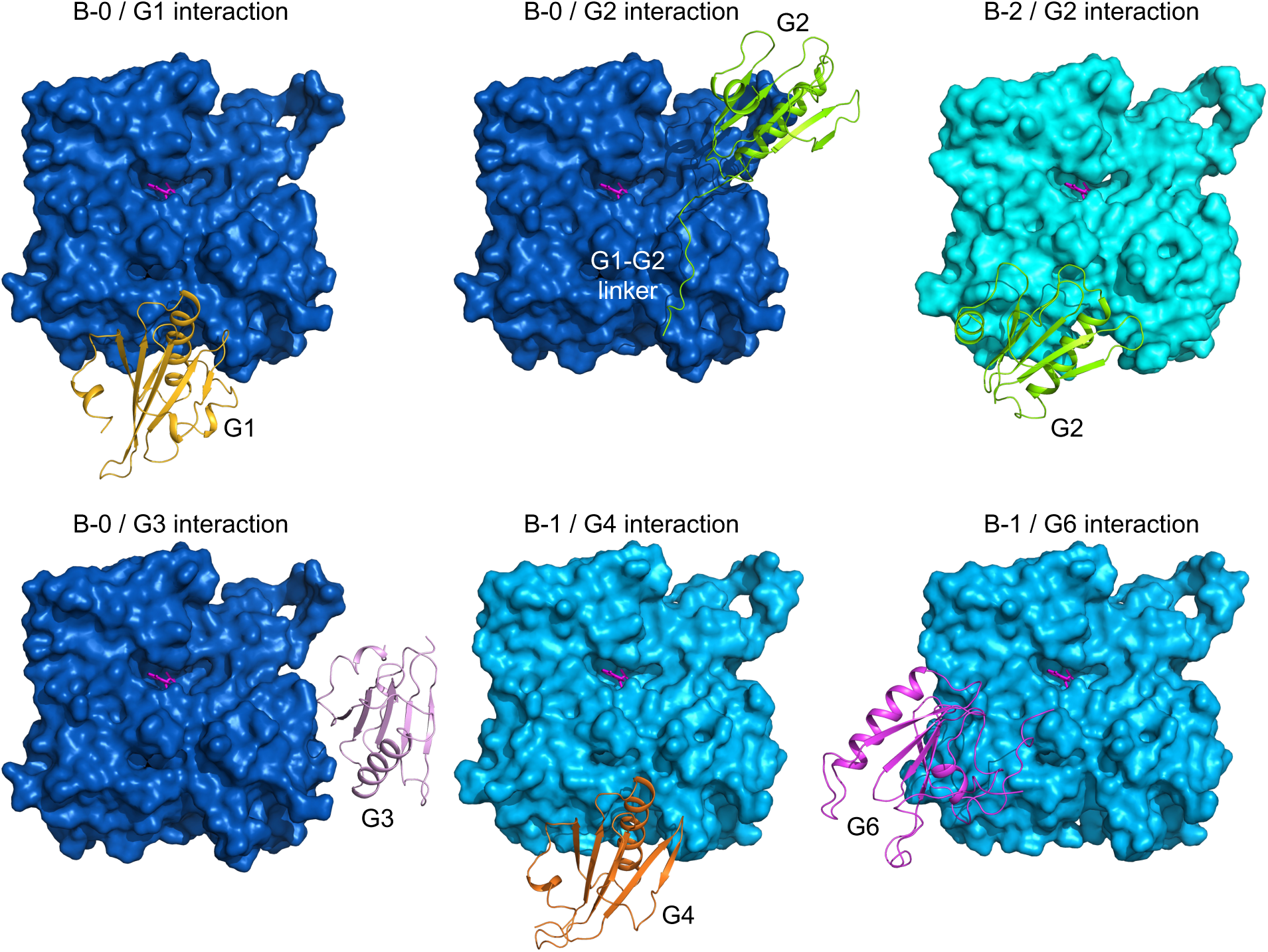
Interactions of actin subunits with gelsolin domains. Interactions of actin subunits with gelsolin domains colored according to Fig. 1b in the main text. Note that three of the gelsolin domains (G1, G2, and G4) bind to a similar area on actin subunits. However, G2 cannot insert its long helix into the hydrophobic cleft of B-2, which is occupied by the D-loop of B-0. Note finally that G5 does not contact any of the actin subunits.

**Extended Data Fig. 10.**
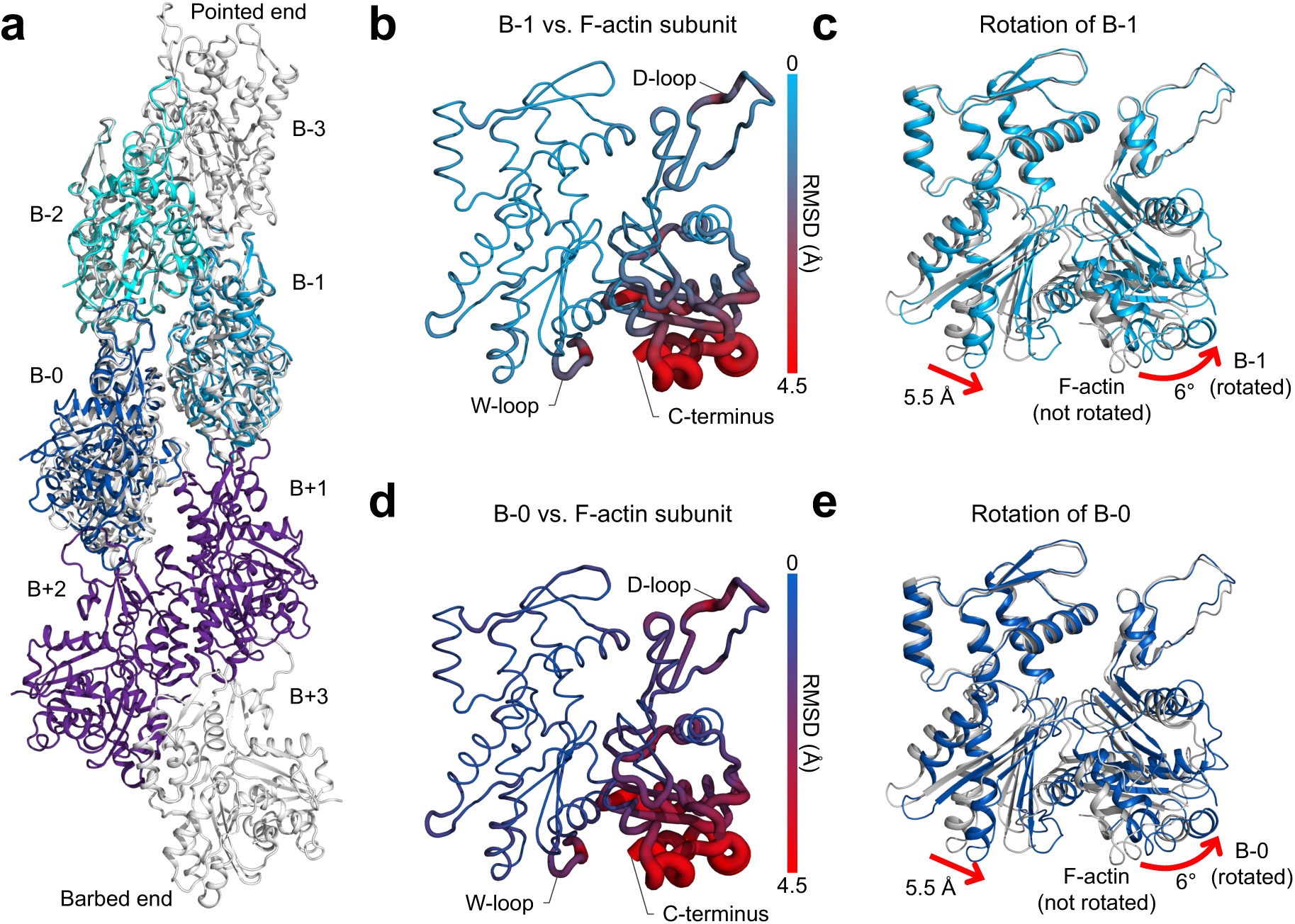
Conformational changes in actin subunits in the G1G3 structure. **a**, Actin subunits B-0, B-1, and B-2 (colored in different shades of blue) from the G1G3-bound structure superimposed onto F-actin (PDB: 8F8P). F-actin subunits are colored grey, except those predicted to interact with G1G3 before severing, which are colored purple. **b** and **d**, Cartoon putty representation of B-1 and B-0 colored and rendered according to RMSD from the corresponding F-actin subunit (as indicated by side bars). Note that the hydrophobic clefts of B-1 and B-0 change substantially due to the binding of G1. The D-loop of these subunits also changes conformation due to interactions with the visible portion of the G3-G4 linker. **c** and **e**, B-1 (light blue) and B-0 (blue) are rotated by 6° compared to their corresponding F-actin subunits (grey), with residues at their barbed ends moving by up to 5.5 Å. Note the rotation of B-1 and B-0 occurs in addition to the changes shown in parts b and d.

